# THE IMPACT OF MISTRANSLATION ON PHENOTYPIC VARIABILITY AND FITNESS

**DOI:** 10.1101/2020.05.19.104141

**Authors:** Laasya Samhita, Parth K Raval, Godwin Stephenson, Shashi Thutupalli, Deepa Agashe

**Affiliations:** National Centre for Biological Sciences, Tata Institute of Fundamental Research, Bangalore, India; International Centre for Theoretical Sciences, Tata Institute of Fundamental Research, Bangalore, India

**Author notes:** Correspondence: Laasya Samhita (,), Deepa Agashe. Equal contribution (Parth K Raval, Godwin Stephenson).

**Keywords:** Mistranslation, non-genetic mechanisms, phenotypic variation, fitness

## Abstract

Phenotypic variation is widespread in natural populations, and can significantly alter their ecology and evolution. Phenotypic variation often reflects underlying genetic variation, but also manifests via non-heritable mechanisms. For instance, translation errors result in about 10% of cellular proteins carrying altered sequences. Thus, proteome diversification arising from translation errors can potentially generate phenotypic variability, in turn increasing variability in the fate of cells or of populations. However, this link remains unverified. We manipulated mistranslation levels in *Escherichia coli*, and measured phenotypic variability between single cells (individual level variation), as well as replicate populations (population level variation). Monitoring growth and survival, we find that mistranslation indeed increases variation across *E. coli* cells, but does not consistently increase variability in growth parameters across replicate populations. Interestingly, although any deviation from the wild type (WT) level of mistranslation reduces fitness in an optimal environment, the increased variation is associated with a survival benefit under stress. Hence, we suggest that mistranslation-induced phenotypic variation can impact growth and survival and has the potential to alter evolutionary trajectories.

## INTRODUCTION

Non-genetic phenotypic variability has long fascinated biologists, not least due to its potential evolutionary impacts. Various aspects of such variation have been analysed from distinct perspectives. Perhaps the best-studied form of non-genetic variation is phenotypic plasticity, when individuals change their phenotype in response to their local environment. Such plasticity – which may be adaptive – is studied largely in animals and plants, and has clear consequences for population as well as community ecology and evolution (reviewed in Bolnick et al. 2011; Raffard et al. 2019). In other cases, only some individuals in a population may respond to environmental change at a given point of time. The resulting heterogeneity in the population potentially represents an evolved bet-hedging strategy, whereby different fractions of the population are better adapted to distinct environments (reviewed in Ackermann 2015). For instance, some cells in *Bacillus subtilis* populations form spores in stressful conditions (Tan and Ramamurthi 2014), whereas others remain metabolically active. Under prolonged stress, the spores stand a better chance of survival; however, if the stress is transient, non-spore formers divide more rapidly. Finally, rather than specific responses to environmental change, genetically identical cells may have distinct phenotypes due to “noise” arising from stochastic variation in gene expression or errors in transcription and translation (Drummond and Wilke 2009; Gout et al. 2013; Ackermann 2015; Carey et al. 2018). Such phenotypic heterogeneity has been well studied in microbial populations, although its evolutionary consequences are relatively poorly understood (Ackermann 2015; van Boxtel et al. 2017).

The evolutionary impacts of non-genetic variation are usually reported either as divergent individual level outcomes (e.g. genetically identical cells with distinct phenotypes may have different reproductive success; Fig. 1a), or as altered mean population-level parameters (e.g. heterogenous populations may grow more slowly than homogeneous populations; Fig. 1b). Both are useful from an evolutionary perspective: growth and survival of individuals ultimately determines trait mean and variance within the population, and hence the outcome of selection. However, non-genetic variation could alter not only average population performance, but also the variance in population level parameters. For example, replicate populations with high individual-level heterogeneity may have more variable average growth rates than replicate homogeneous populations (Fig. 1c). This may occur, for example, due to stochastic effects in each replicate, or due to divergent outcomes of interactions between individuals in each population. Thus, despite similar initial heterogeneity within each replicate, the population level outcomes may be divergent (right hand plot in Fig. 1c). Such effects on variance are important to measure, because higher variance in evolutionary outcomes between populations reduces the repeatability (and hence predictability) of evolutionary dynamics. However, the potential impact of between-individual phenotypic heterogeneity on the variance in population level parameters remains unexplored. Hence, it is unclear whether a mechanism that generates individual level variation (i.e. between cells or organisms) also consistently generates divergent population-level outcomes.

**Figure 1.**
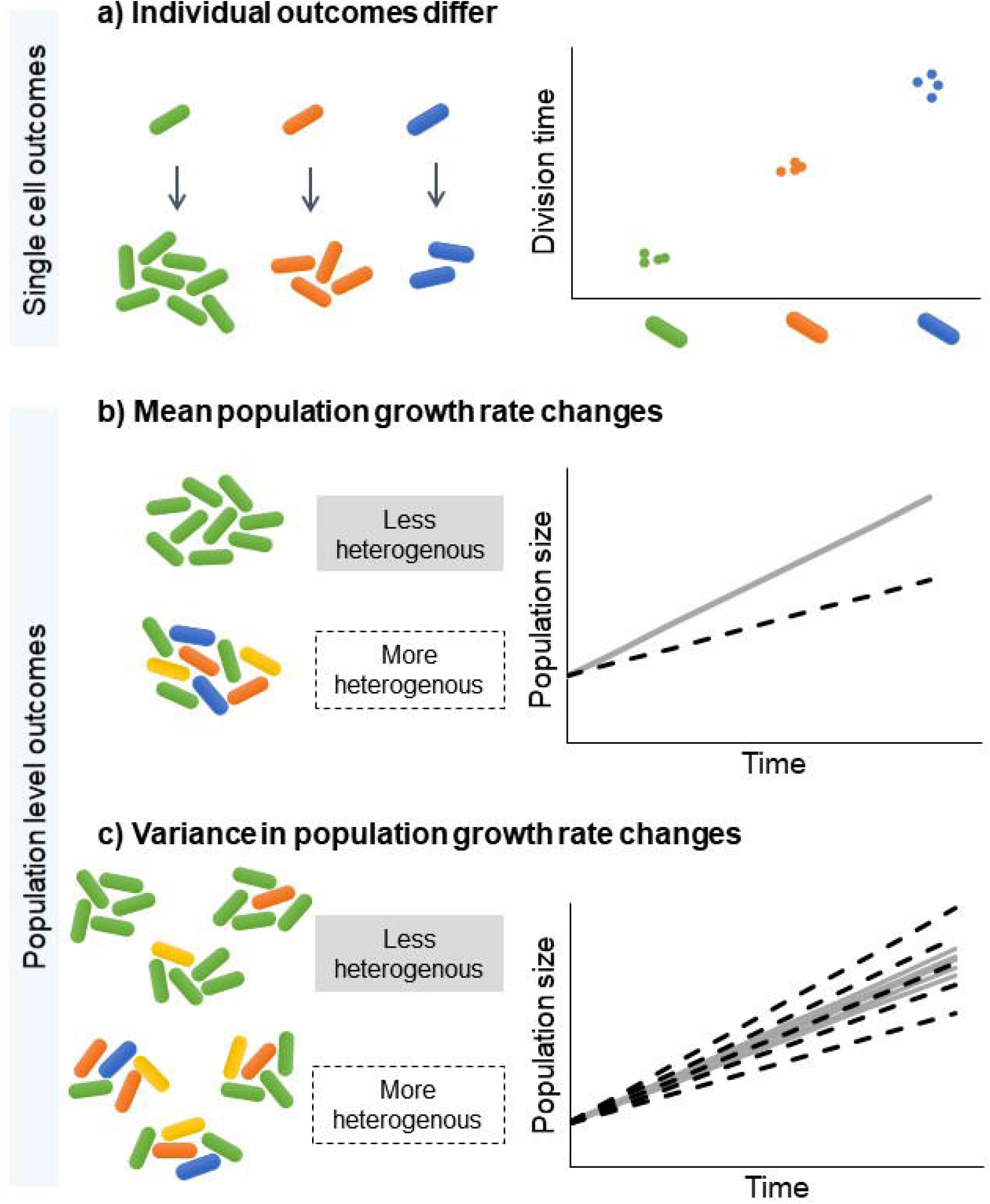
Possible impacts of individual or cell-to-cell heterogeneity on individuals and populations. (a) Cell to cell heterogeneity (indicated by cell colour) causes variation in individual fitness parameters such as the rate of cell division. (b) Populations that have a large amount of cell to cell heterogeneity may grow more slowly, because of the phenotypic load generated by slower-dividing cells. (c) Cell to cell heterogeneity can impact between-population variance due to stochastic effects or diverse cell to cell interactions. As a result, replicate populations with high individual-level heterogeneity may show more variable growth rates (spread around the mean value) than replicate homogeneous populations.

From a mechanistic perspective, translation errors are especially interesting because they are an inescapable aspect of the biology of all life forms, and they occur at a high rate. For instance, in *Escherichia coli,* about 10% of dihydrofolate reductase enzyme molecules differ from the native sequence of the protein (Ruan et al. 2008). The typical mistranslation rate is ∼1 in 10^4^ incorrect amino acids in a growing protein chain, increasing to as high as 1 in 10^3^ amino acids under stress (Ribas de Pouplana et al. 2014; Mordret et al. 2018). Such high error rates can generate significant proteome diversity (Nakahigashi et al. 2016; Mordret et al. 2018). Importantly, unlike many other forms of non-genetic variability in microbes – such as spore formation and persister cells (Ackerman 2015) – mistranslation can generate continuous (rather than binary) phenotypic variation, allowing a more fine-tuned response to a large diversity of stresses. Such continuous variation in protein quality or quantity can reliably generate large phenotypic variability, as seen with the heat shock protein Hsp90 (Cowen and Lindquist 2005) and prions (Halfmann et al. 2012), which in turn can determine survival in a new environment (Novick and Weiner 1957). Therefore, it is speculated that mistranslation-induced non-genetic variation may generate substantial phenotypic variability, potentially altering the outcome of natural selection (Miranda et al. 2013; van Boxtel et al. 2017).

However, postulating a general hypothesis about the evolutionary consequences of mistranslation-induced variation requires consideration of multiple nuances. First, proteome diversity is visible to natural selection only if it leads to phenotypic diversity in traits that influence fitness. Given various buffering mechanisms driven by chaperones and the degradation of mistranslated products (Bratulic et al. 2015; Kalapis et al. 2015), protein diversity may not always generate phenotypic diversity. Hence, in the absence of this link, proteome diversity is of little evolutionary consequence. Second, the effects of mistranslation are inherently unpredictable and not heritable, weakening the potential for long-term consequences. In microbes such as *E. coli,* proteome diversity has limited across-generation persistence due to protein dilution at cell division. Hence, favourable mistranslated protein variants may never be sampled again, limiting their effect on evolutionary dynamics. Finally, in a constant optimal environment, populations should face stabilizing selection. This means that any mechanism that generates increased variability between individuals is likely to move them away from the optimal phenotype, creating a ‘phenotypic load’. Therefore, if mistranslation increases cell to cell variability, it is likely to be adaptive primarily under directional or disruptive selection, such as might be imposed in a new environment or under stress. In contrast, in a constant environment, mistranslation is more likely to be maladaptive. These limitations of the evolutionary consequences of mistranslation remain largely untested. Previous work shows that increased mistranslation can generate diversity in cell morphology and cell surface receptors (Bezerra et al. 2013; Miranda et al. 2013). However, there is limited experimental evidence directly linking mistranslation with phenotypic variation relevant to fitness.

Here, we tested whether altering mistranslation levels in *E. coli* impacts variability at both single cell and population levels, in phenotypes relevant for growth and survival. We increased the basal level of mistranslation in wild type (WT) cells by introducing mutations or by changing the growth environment. Recently, we showed that generalised mistranslation increases mean population survival under specific stresses (Samhita et al. 2020). However, we had not explored whether mistranslation generates phenotypic variability that influences both cell and population fitness, and in optimal as well as stressful environments. Here, we find that mistranslation indeed increases phenotypic diversity in *E. coli* at the single cell level, and that suppressing mistranslation via hyper-accurate ribosomes reduces this variability. However, increased single-cell variability did not affect variability in population level growth parameters. Importantly, while mistranslation-associated variability is costly in optimal conditions, it increases survival under stress; and this effect is observed even with a transient increase in mistranslation. Thus, mistranslation indeed results in phenotypic diversification across cells, and this diversity is directly correlated with survival under stress.

## RESULTS

### Mistranslation increases cell-to-cell but not between-population variability in growth and division time

To test the impact of mistranslation on phenotypic variation, we manipulated basal mistranslation levels and measured division time and cell length of GFP-tagged single cells in a microfluidics device (Fig. 2a). Cell division time is a key proxy for fitness under optimal growth conditions, and changes in cell length are predictive of the physiological state of a cell (Wehrens et al. 2018). We genetically increased mistranslation levels in our WT *E. coli*, generating the “Mutant”: a strain with depleted initiator tRNA content that has increased mistranslation rate (Samhita et al. 2013). Conversely, we reduced mistranslation rate by introducing a mutation in the ribosomal protein S12. Other than these genetic manipulations, we also increased mistranslation rates by adding the amino acid analogues canavanine or norleucine, or the antibiotic streptomycin to the WT growth media. In each case, we measured phenotypic variability across cells and across populations. Because the data were not normally distributed, we compared distributions for spread around the median using the Fligner-Killeen test.

**Figure 2.**
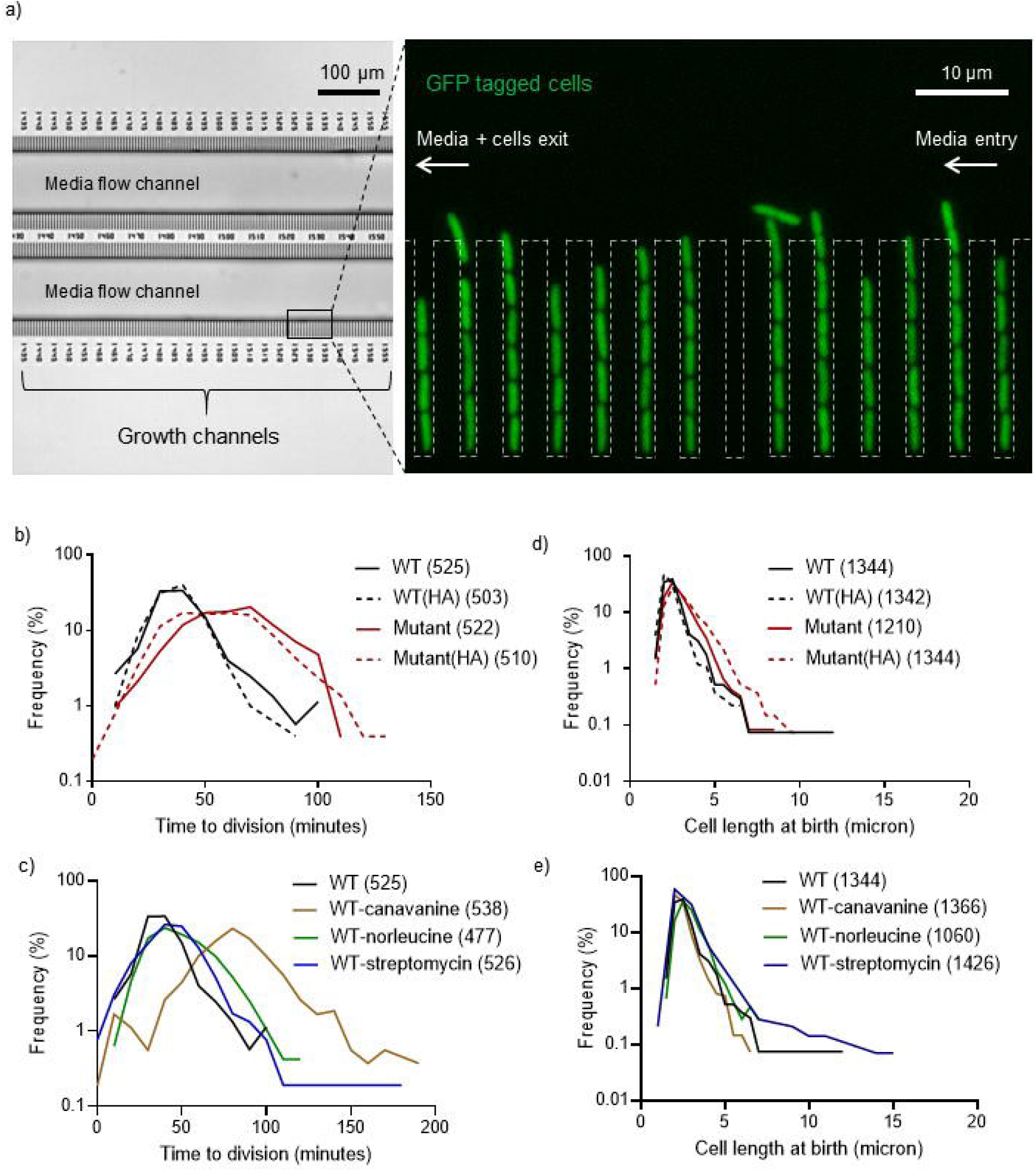
Mistranslation increases cell-to-cell variability in growth and division time: We injected ∼10^5^ cells of the indicated strains into a microfluidic device designed for single cell tracking, and monitored cell growth and cell length under the microscope. (a) Schematic of the microfluidics device showing GFP tagged *E. coli* single cells growing and dividing in channels within the device (b) – (e) Frequency distributions of cell length and division time of single cells as monitored in the microfluidics device. For (b) and (d), the center of each bin (class interval) is 5 units; for (c) and (e) 0.5 units; and each such point is connected to the next one. Wider distributions indicate greater cell-to-cell variation. Total number of cells (n) is indicated within parentheses in the key. WT=wild type; HA=hyper-accurate.

As predicted, all methods of increasing mistranslation increased cell-to-cell variation in the time to division (Fig. 2b–c; Fligner Killeen test: Mutant >WT, Χ^2^=9, P < 0.0001; WT_can_>WT, Χ^2^=84.9, P < 0.0001;WT_nor_>WT, Χ^2^=54.1, P < 0.0001; WT_strp_>WT, Χ^2^=32.6, P < 0.0001;Table S1) and most led to increased cell size variability (Fig. 2d–e; Fligner Killeen test: Mutant>WT, Χ^2^=86.9, P < 0.0001; WT_can_<WT, Χ =46.4, P < 0.0001; WT_nor_>WT, Χ^2^=58.5, P < 0.0001; >WT vs WT_strp_, ns, Χ^2^=1.5, P =0.2; Table S1). Conversely, reducing mistranslation via hyper-accurate ribosomes reduced variability in cell size but not division time of the WT, but did not reduce either for the Mutant (Fig. 2b and 2d; See Table S1). In the analyses described above, for each strain we pooled data across ∼60 channels of the microfluidics device (each with a single, original mother cell) and ∼600 divisions. To confirm that this pooling did not end up averaging differences across generations, we re-analysed data focusing only on the first three divisions of each mother cell (∼60 cells in total); and found similar results (Fig. S2). Thus, our microfluidics experiments show that mistranslation directly increases single cell phenotypic variability in growth and division time.

Next, we examined the consequences of mistranslation on variability at the population level. We compared growth curves of ∼40 replicate populations per strain/condition (Fig. S3), to test whether cell to cell variability manifests at the population level in growth parameters of mistranslating strains. Interestingly, in most cases increasing mistranslation did not increase between-population variability in growth rate, lag time (time until culture reaches OD_600_ ∼0.02), or growth yield (Fig. 3; Table S1). The only exceptions were increased between-population variability in WT growth rate due to streptomycin (Fig. 3a), and WT yield due to streptomycin or norleucine (Fig. 3b; Table S1). Contrary to expectation, reducing mistranslation through hyper-accurate ribosomes also increased variability in the WT lag time and in all three parameters for the Mutant (Fig. 3; Table S1). Thus, in contrast to single-cell variability, mistranslation did not consistently affect variability in population level growth parameters.

**Figure 3.**
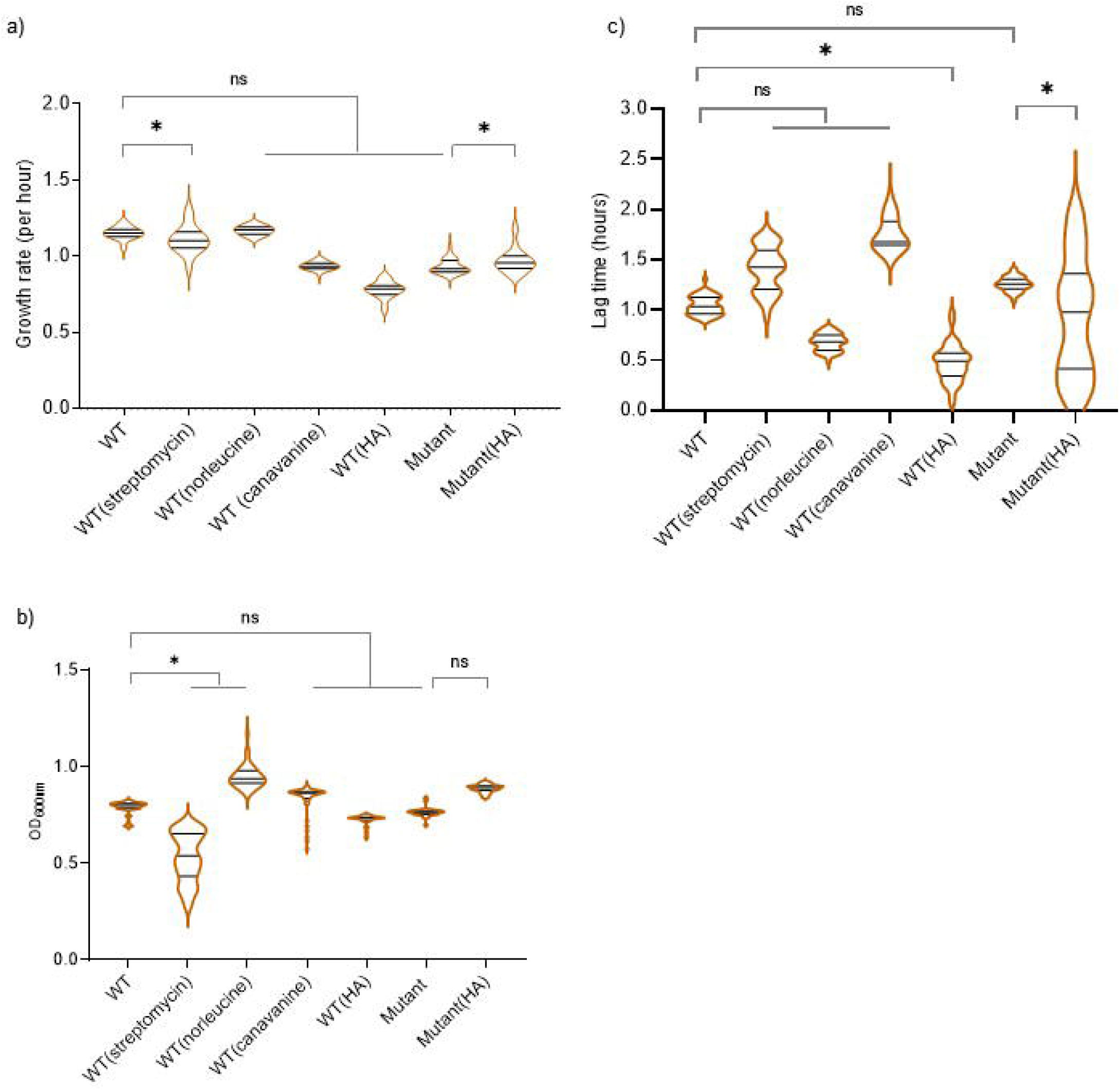
Mistranslation does not impact variability in growth parameters across replicate populations: Violin plots showing the distribution of three population growth parameters, estimated using ∼40 (37 to 44) biological replicates (populations) for each strain or growth condition. (a–b) Growth rate (c–d) growth yield and (e–f) lag time (time until culture reaches OD_600_ ∼0.02). Median, 25^th^ and 75^th^ quartiles are indicated by solid lines within each violin. The length of each violin corresponds to the range of the distribution. Asterisks indicate significant differences in variability. WT=wild type; HA=hyper-accurate.

### Mistranslation is costly under normal conditions

In addition to affecting variability in single cell growth parameters, altering mistranslation levels often incurred a cost. At the single cell level, median time to division increased significantly with higher mistranslation, although reducing mistranslation had no effect (e.g. WT: 36 min, Mutant: 62 min; Mann-Whitney test, U=45832, P<0.001; Fig. 2b and 2c; Table S1). Increased mistranslation also increased cell length in most cases (Fig 2d and 2e; WT 2.4 μm vs Mutant 2.7 μm, Mann-Whitney test, U=586570, P<0.0001; other comparisons in Table S1), whereas reducing WT mistranslation through hyper-accurate ribosomes decreased cell length (WT 2.4 vs WT(HA) 2.2 μm, Mann-Whitney test, U=710144, P<0.001). Increased cell length might partly account for greater division times of mistranslating strains, although we did not explicitly test this. Note that while increased cell length is associated with stressful conditions (Wehrens et al. 2018), it is not clear if longer cells are necessarily a cost here, given the associated increase in biomass. These patterns at the single cell level were also reflected in population level parameters. Populations with either increased or decreased mistranslation relative to WT had lower growth rate and greater lag time for the most part, although growth yield did not change consistently (compare median values in Fig. 3a and 3c; also see Fig. S4 and Table S1). Overall, mistranslating cells were longer and divided more slowly than the WT, and mistranslating populations showed slower growth; suggesting a cost of mistranslation.

### Mistranslation increases population survival under stress

Although costly under normal conditions, previous studies suggest that mistranslation often confers a benefit under stress at the population level (reviewed in Mohler and Ibba 2017). We therefore examined the impact of two stresses – high temperature (42°C) and starvation (Koch 1971; van Elsas et al. 2011) – on single cell and on population growth parameters. Both WT and Mutant single cells divided faster at 42°C than at 37°C, but the increase was much larger in the Mutant (compare Fig. 4a vs. Fig. 2b; median division time WT (42°C): 28 min vs. WT (37°C): 36 min, Mann-Whitney test, U=89652, P<0.0001; Mutant (42°C): 38 min vs. Mutant (37°C): 62 min, U=66956, P<0.0001). Thus, the cost of mistranslation decreased at high temperature; but the Mutant still took longer to divide than the WT (median division time: Mutant 38 min vs. WT 28 min, Mann-Whitney test, U=123307, P<0.0001). At 42°C, reducing mistranslation rate was also slightly costly for the Mutant (median division time: Mutant (HA) 40 min, Mutant 38 min, Mann-Whitney test, U=149832, P=0.003), and more so for the WT (median division time WT(HA) 34 min >WT 28 min, Mann-Whitney test, U=118457, P<0.0001) (Fig. 4a). Overall, an increase in temperature reduced the cost of slow growth and mistranslation (as measured by division time difference) in the Mutant, but did not give it a growth advantage over the WT. All else being equal, division time is a good measure of fitness in actively dividing cells; but once cells enter stationary phase, division rate decreases and growth rate no longer determines competitive fitness. We therefore examined longer term survival as a population level fitness measure, assessing total viable counts in 48 h (stationary phase) cultures expose to high temperature. Across populations, mutant survivability was higher than WT at both 37°C and 42°C, with a stronger effect at 42°C (Fig. 4b; Table S1). Thus, mistranslation was either beneficial or less costly for cells and populations exposed to high temperature stress.

**Figure 4.**
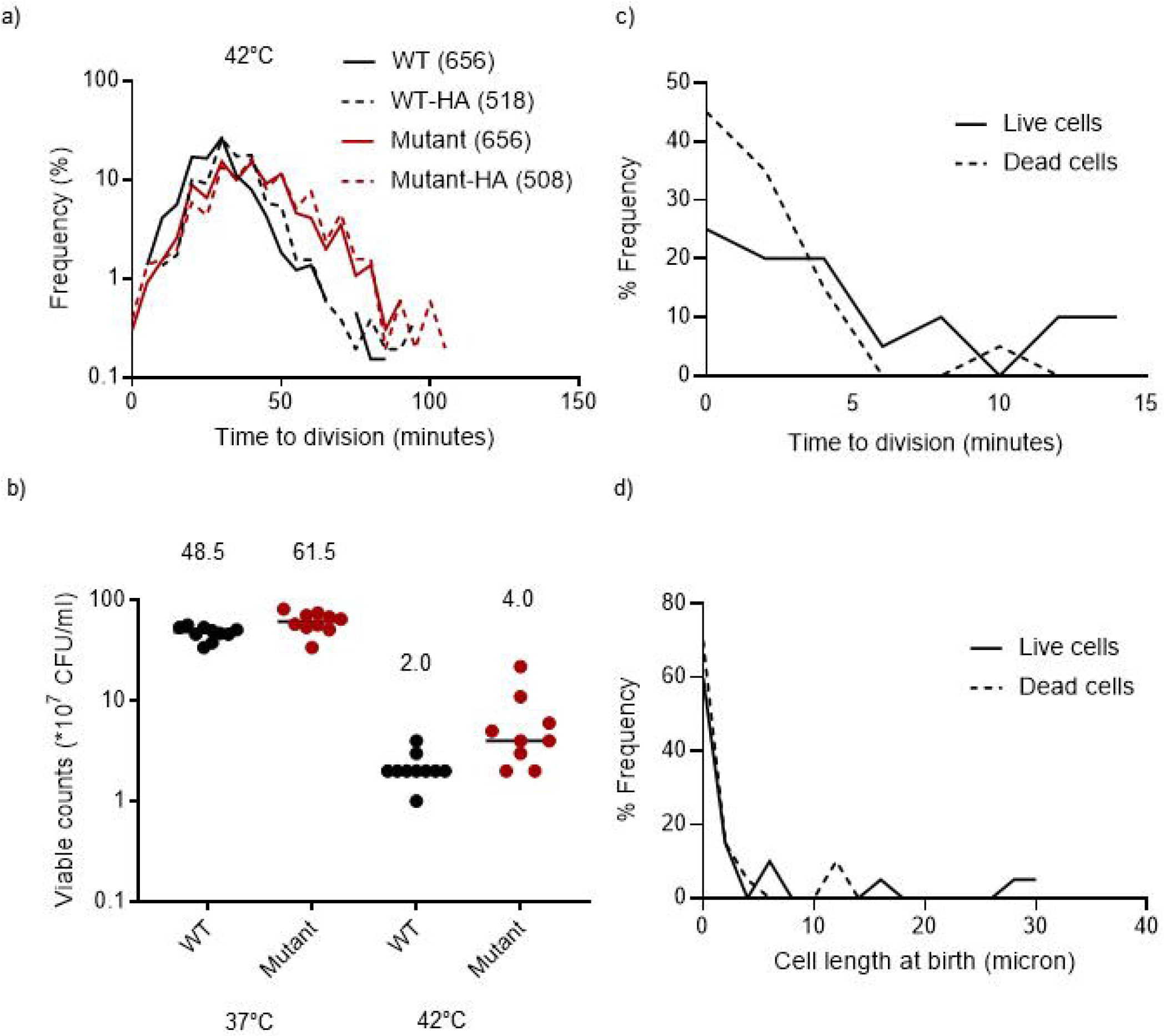
Mistranslation increases survival under stress: (a) Frequency distributions of division time of single cells of different strains monitored in the microfluidics device at 42°C. The center of each bin (class interval) is 5 units, and each such point is connected to the next one by a line. WT=wild type; HA=hyper-accurate. (b) Total viable counts (estimated by dilution plating) in WT and Mutant cultures (n=9) grown for 48 h (late stationary phase) at 37°C or 42°C. Numbers indicate median viable count in each case. (c–d) Distribution of division time and cell length of single WT cells that subsequently either survived (live cells, n=86) or died (dead cells, n=31) after being starved of nutrients for ∼10 h, in saline. Overall, WT cells had a higher fraction of dead cells than the Mutant (see Results).

Next, we tested cell survival under starvation stress. We introduced diluted overnight cultures in the microfluidics device, allowed them to enter mid-log phase and then added saline (0.85% NaCl) instead of growth medium, to test whether Mutant and WT had different rates of cell death in the absence of nutrients (where no cell division can occur). We did this across three different experimental blocks, sampling ∼650 cells of WT and Mutant each, in total (Fig. S5). After 10 h, repeatably, on average ∼19% WT cells died, while only ∼4% Mutant cells died; suggesting that the Mutant is more robust to a lack of nutrients. To test whether dead cells were more likely to have a specific phenotype, we retrospectively measured the time to division and cell length of 31 dead cells and 86 live cells of WT from one set.

Interestingly, both surviving and dead cells were drawn from across the original distribution of cell length and division time (Fig. 4 c–d), indicating that survival was not linked to these specific aspects of cellular level heterogeneity. We could not do a similar analysis for the Mutant, due to the small number of dead cells. Our results suggest that mistranslation increases cell survival under starvation-induced stress; but that the survival is not directly connected to the indicators of cellular phenotype that we measured.

Lastly, we assessed the impact of mistranslation on population survival using competitive fitness. We allowed WT to compete with its hyper-accurate derivative or with the Mutant, using actively growing (log phase) or stationary phase cultures (under starvation) as a starting point. As expected from their relative log phase growth rates (Fig. 3a), WT outcompeted both the mutant and the hyper-accurate strains (Fig. 5a–b and Fig. S6). However, when competing in stationary phase, Mutant had comparable or marginally higher fitness than the WT, both at 37°C and 42°C (Fig. 5c–d). Thus, both increasing or decreasing mistranslation levels in the WT imposed a fitness cost in nutrient rich conditions when rapid growth is favoured. In contrast, under stress, cells with higher mistranslation rates could either co-exist with or perform slightly better than cells with a lower mistranslation rate. Together, our results indicate that mistranslation is costly for growth under optimal conditions, but is often beneficial for survival under stress, at the single cell as well as population levels.

**Figure 5.**
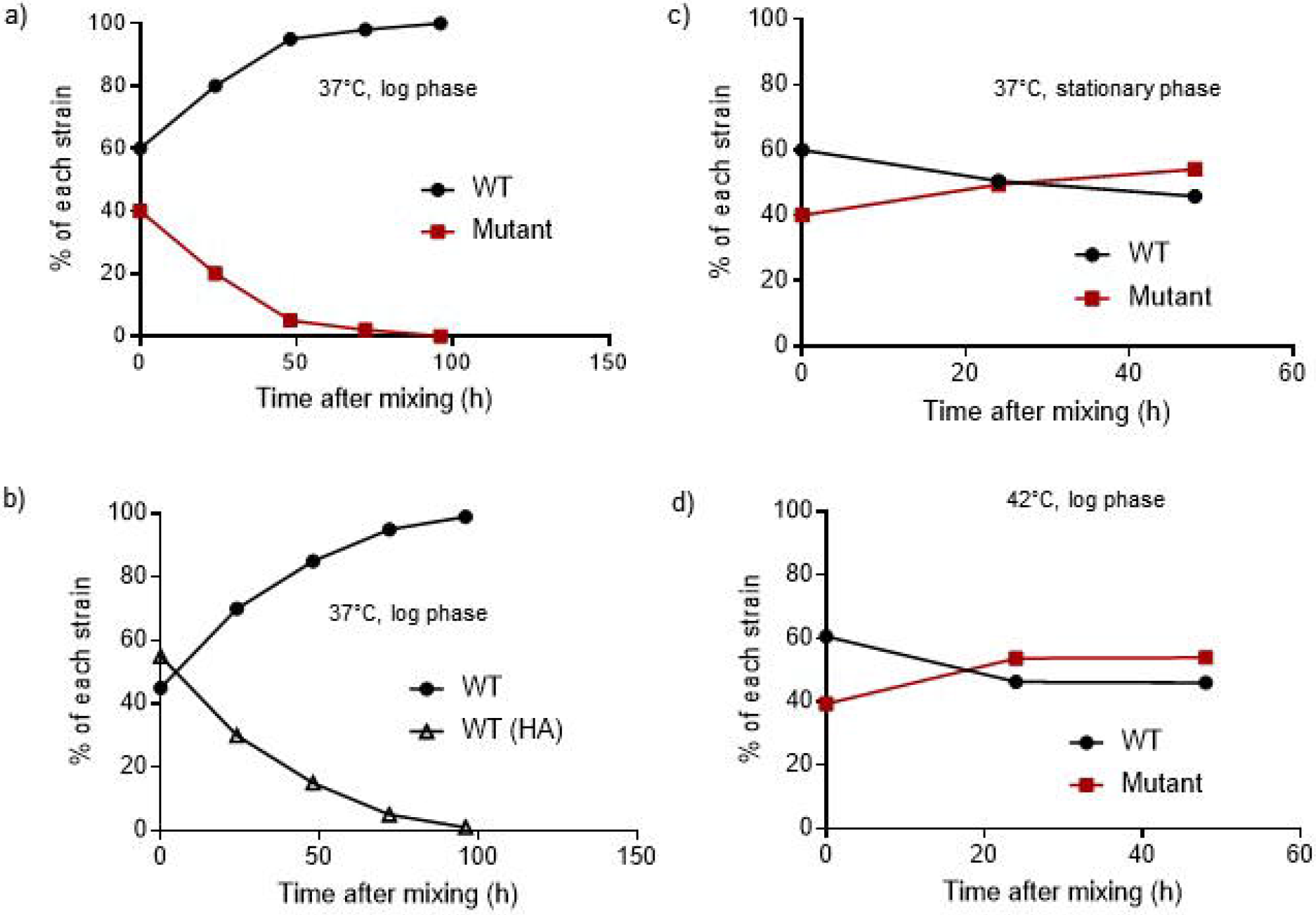
Mistranslation is costly under optimal conditions but beneficial under stress: (a–b) Cell survival as a function of time, during pairwise competition in the log phase of growth at 37°C. We mixed log phase cultures (OD_600_∼0.6) of two strains (as indicated) in LB, and plated aliquots on MacConkey’s agar to estimate survival of each strain. Data for a representative set for each case are shown here; other sets are shown in Fig. S6 (c) Cell survival as a function of time, during pairwise competition in the stationary phase of growth at 37°C. We allowed WT and Mutant cultures to grow independently for 48 h in LB medium and then mixed them to assess competition in late stationary phase. Data for a representative set for each case are shown here; other sets are shown in Fig. S6 (d) Cell survival as a function of time, during pairwise competition in the log phase of growth at 42°C.

### The degree of mistranslation does not correlate with between-population variability

While single cell variability clearly increased with mistranslation, population level variability did not show a clear correlation. To tease apart the role of mistranslation in generating population variability, we exposed WT cells to a gradient of mistranslation, by treating them with increasing concentrations of mistranslating agents (canavanine, norleucine or streptomycin). We expected that across-replicate variability in population growth rate, yield and lag time (estimated using the inter-quartile range of) would increase monotonically, as a result of increasing mistranslation. However, we found that the degree of mistranslation was not strongly correlated with either the median trait values (Fig. 6a-c) or the variability across replicates (Fig. 6d-f). Furthermore, the patterns varied across mistranslating agents, potentially driven by the specific mode of action of each agent, or its impact on other cellular processes unrelated to mistranslation (see Discussion).

**Figure 6.**
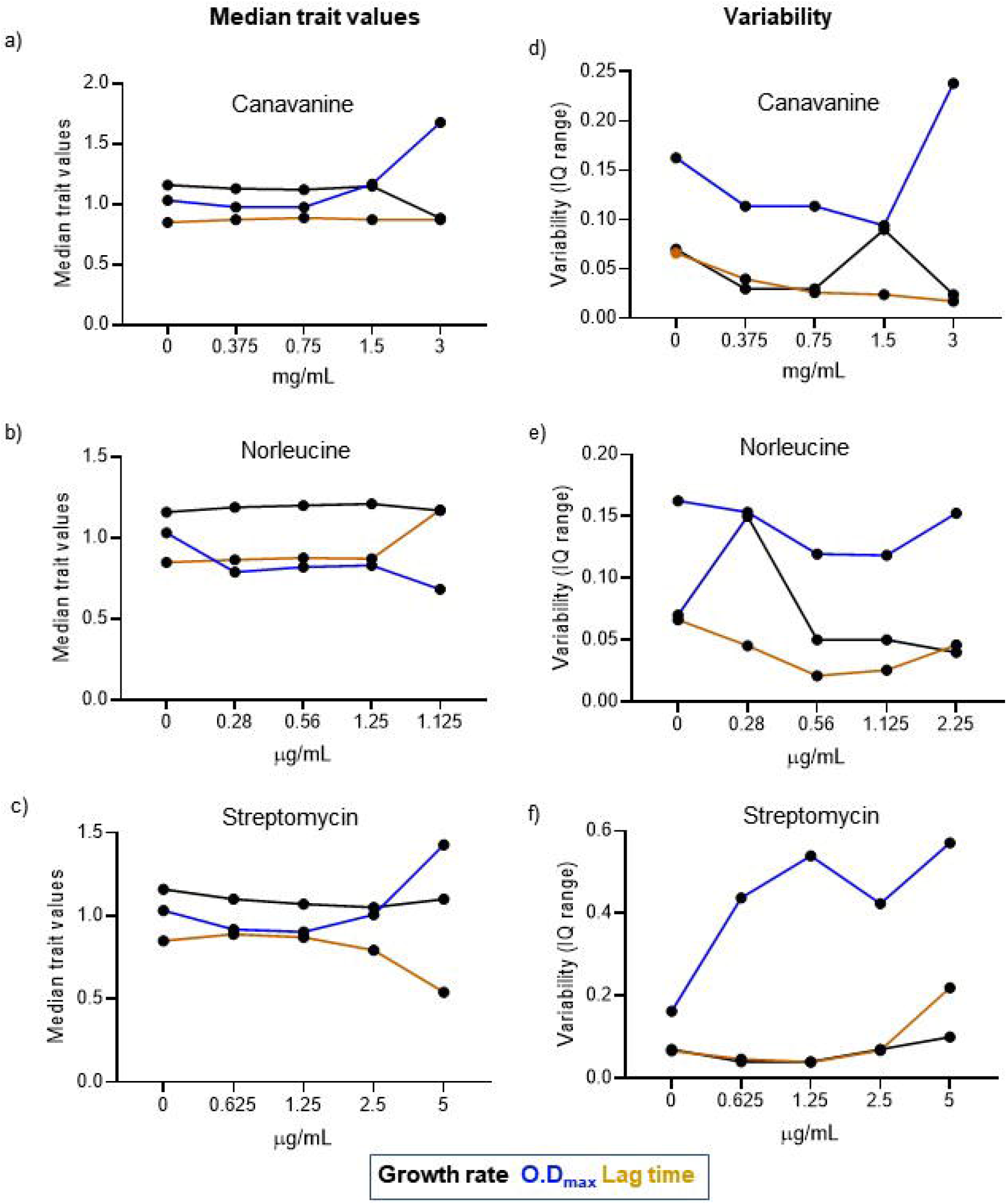
Variability in growth parameters across populations is not correlated with degree of mistranslation: (a–c) Median values for population growth rate, growth yield (OD_max600_) and lag time (time until culture reaches OD_600_ ∼0.02) as estimated from raw growth curves for WT populations treated with four concentrations of canavanine, norleucine or streptomycin (n=∼40, 37 to 44 per treatment). Untreated WT is indicated as zero concentration in all treatments. (d–f) Variability in population growth parameters across biological replicates (n=∼40), estimated using the difference between the 75^th^ and 25^th^ percentile values for each parameter. Untreated WT is indicated as zero concentration in all treatments. Linear regression for variability in growth rate as a function of canavanine, norleucine or streptomycin concentration: R^2^=0.5, 0.3 and 0.3; for OD_max_: R^2^=0.8, 0.28 and 0.48; for lag time: R^2^= 0.02, 0 and 0.6 respectively.

### A brief burst of mistranslation is sufficient to increase subsequent population survival

In the experiments described so far, we maintained a constant level of mistranslation throughout the course of the experiment, because mistranslation generates fresh phenotypic variability in each generation. As discussed in the Introduction, without such renewal, the impact of initial mistranslation should diminish over successive generations. However, proteome changes can be transferred across generations in other ways, such as through protein aggregates (Govers et al. 2018). We therefore asked whether a brief pulse of mistranslation can alter subsequent cell viability, and whether this effect scales with the degree of initial mistranslation, both under normal growth conditions and under stress (high temperature). We kept the window of exposure to the stress to within 2 doubling times of the slowest strain (∼2 h), so that any effects we observed were solely due to the mistranslating agent and not confounded by subsequent selection on survivors. We also ensured that the concentrations of mistranslating agents and the magnitude of the stress used did not cause any cell death within this window.

We found that briefly exposing cells to increasing concentrations of canavanine, norleucine or streptomycin increased survival on LB agar at 37°C (Fig. 7a), consistent with our prior observations with the Mutant (Fig. 4b and Fig. 5c-d). Streptomycin was an interesting outlier, with an intermediate concentration consistently maximizing survival. We speculate that this concentration indicates a threshold beyond which the toxic effects of mistranslation overwhelm its benefits. With increasing concentrations of mistranslating agents, as before (Fig. 6a-c), we did not find a consistent trend towards higher median survival, except with streptomycin (linear regression: WT_nor_, R^2^=0.005, P=0.4; WT_can_, R^2^=0.05, P=0.18; WT_strp_, R^2^=0.4, P=0.0004). Surprisingly, at 42°C, none of the treatments significantly increased mean survival as compared with the WT (Fig. 7b). Again, survival increased with increasing concentration of streptomycin, but the other mistranslating agents did not have this effect (linear regression: WT_nor_, R^2^=0.001, P=0.9; WT_can_, R^2^=0.03, P=0.6; WT_strp_, R^2^=0.6, P=0.02). Part of the reason for the lack of difference at 42°C could be the small sample size. For logistical reasons (see Methods), we could not increase our sample size at 42°C; given the large variability across replicates, we may thus have limited power to detect a correlation.

**Figure 7.**
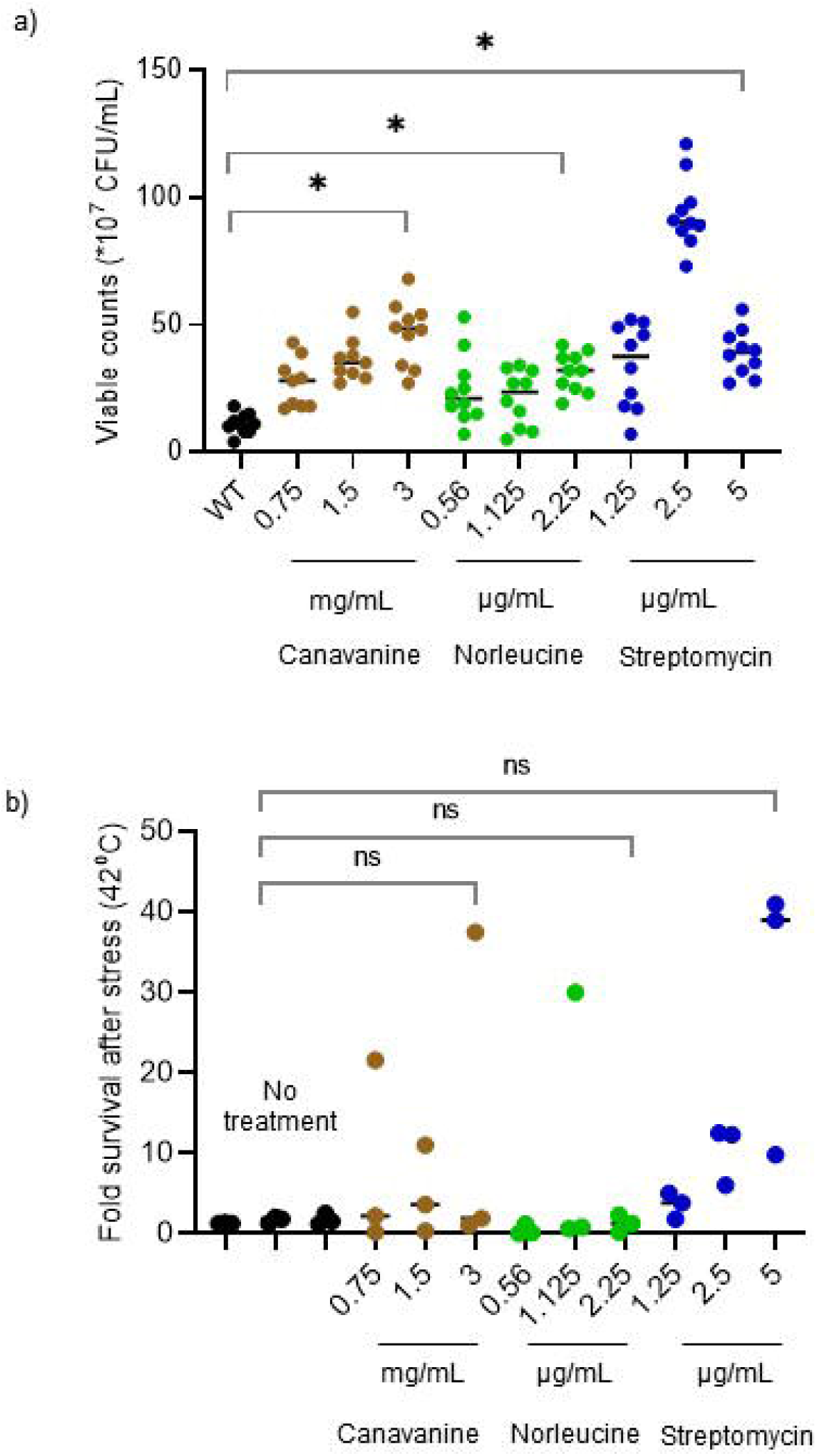
A brief burst of mistranslation enhances survival under optimal conditions. a) Effect of treatment with different concentrations of canavanine, norleucine or streptomycin on viable cell counts at 37°C (n=10 per concentration). We treated log phase cultures (OD_600_∼0.6) of each strain indicated with three concentrations of each mistranslating agent as indicated, for 2 h. Cells were then spun down, washed and dilution plated followed by incubation at 37°C for 24 h. (b) Effect of a brief exposure to different concentrations of canavanine, norleucine or streptomycin on cell survival at 42°C (n=3 per concentration). We treated cells as above; then spun down, washed, and incubated them at 42°C for a further 2 hours in fresh medium. To calculate fold change in survival due to high temperature, we dilution plated and incubated cells at 37°C for 24 h. Horizontal bars indicate median values in both panels. Asterisks indicate significant differences in the median values.

Overall, as with population growth parameters (Fig. 6), there was no clear correlation between the extent of mistranslation and magnitude of the survival benefit at the population level. However, a brief increase in mistranslation did increase subsequent survival. In conjunction with previous results (Fig. 4b and Fig. 5c-d), these results suggest that mistranslation has the potential to influence longer-term population and evolutionary dynamics.

## DISCUSSION

Translational errors have been intensively studied by molecular biologists, leading to a detailed understanding of their causes and immediate cellular consequences (Kramer and Farabaugh 2007; reviewed in Ribas de Pouplana et al. 2014). At the same time, evolutionary biologists have analysed the broader consequences of errors in cellular processes for non-genetic adaptation (Whitehead et al. 2008; Evans et al. 2018). However, these two perspectives have only rarely been connected, resulting in poor empirical understanding of the possible role of mistranslation for survival and adaptation in new environments. By directly manipulating mistranslation rates in multiple ways, we provide clear empirical evidence that mistranslation introduces phenotypic variability across cells; but does not consistently introduce variability across populations. Furthermore, cell-to-cell variability has environment-dependent impacts on fitness: altering WT mistranslation rates in either direction is deleterious under optimal conditions, whereas mistranslation-induced variation is associated with improved survival under stress (Fig. 8). Recently, using the same manipulations, we showed that global mistranslation increases survival under at least two stresses, DNA damage and high temperature (Samhita et al. 2020). Together, these results show that mistranslation-induced variability has the potential to significantly alter ecological and evolutionary dynamics of populations.

**Figure 8.**
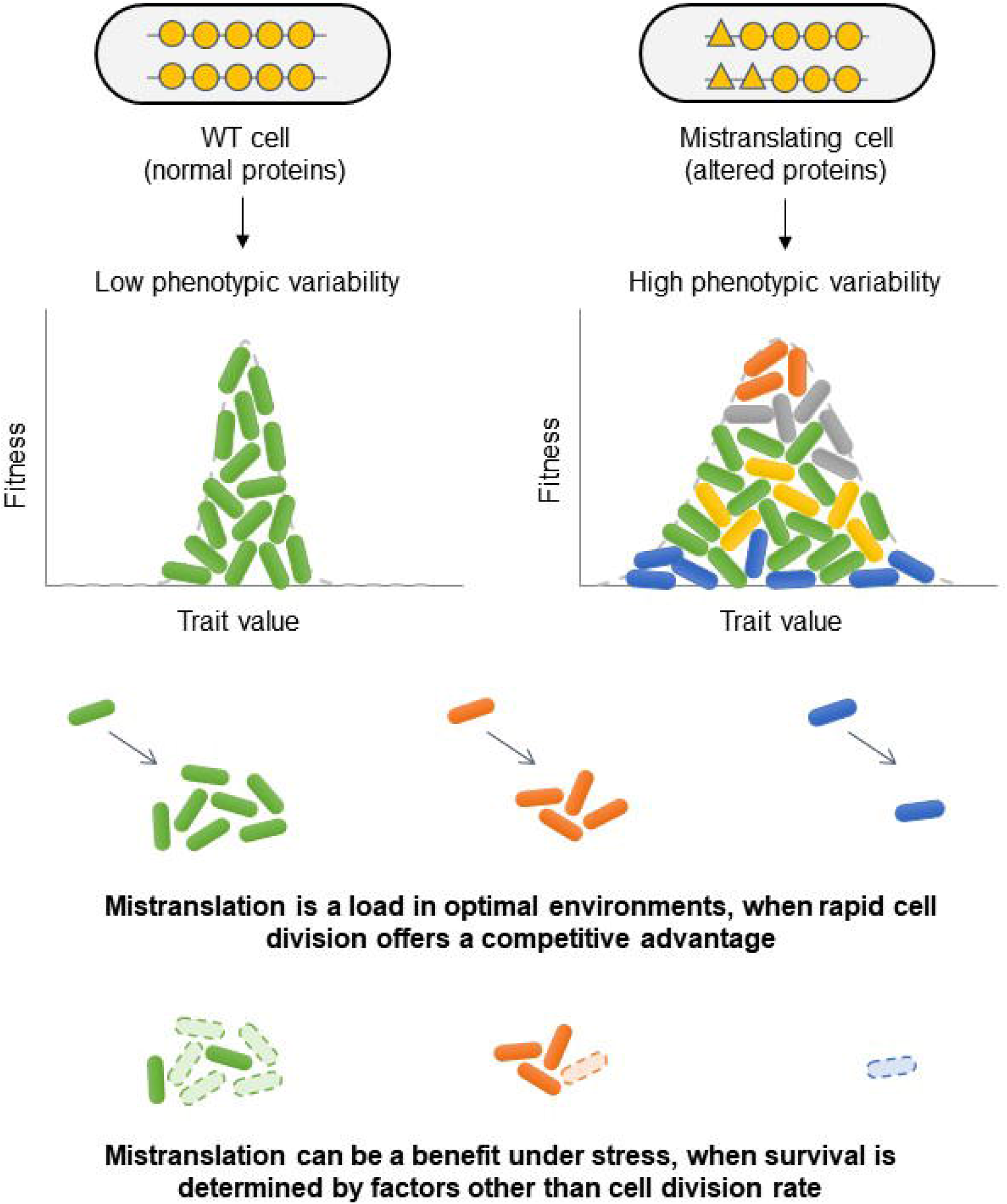
Summary for the impact of mistranslation-induced variability on fitness.

Note that although mistranslation generated variability, it was difficult to establish whether the variability itself was directly responsible for growth and survival benefits; or whether these benefits arose due to secondary effects of mistranslation (see discussion below). It is also tempting to speculate that the increased variability leads to a bet hedging strategy, i.e. a specific sub-population within a heterogeneous population (for example, cells with the highest growth rates, at one end of a long tail in the distribution) performs best in a new environment. If so, the parent (non-diversified) population that lacks this subpopulation would be at a disadvantage, leading to eventual over-representation of the subpopulation. However, our single cell experiments do not support such a bet hedging strategy within the fitness parameters that we measured: cells that survived starvation were not drawn from a specific region of the overall distribution of cell size and time to division (Fig. 4c-d). It is still possible that mistranslating cells do sample a specific sub-population and exhibit bet hedging, with reference to some other phenotype that we have not tested. For example, in the expression level of a specific protein. Further experiments are necessary to test whether (and under what conditions) mistranslation-induced single-cell variability may serve as a general bet-hedging strategy, or consistently increase population growth or survival.

Interestingly, we did not find strong support for our hypothesis that within-population variability can also generate across-population variability. Thus, despite the significant between-cell variation introduced by mistranslation, population behaviour largely remains repeatable (and hence predictable). What explains this discrepancy? It is possible that averaging across several single cell distributions may hide the underlying variability. For example in our case, mistranslating cells have more variable division times. However, if all populations consistently generate the same set of alternate proteomes (or distribution of division times), then we would not see significant variation across replicate populations. Alternatively, selection during the population growth cycle may ensure that only the fastest-growing cells contribute to population growth rate, reducing the magnitude of variation across replicate populations. This is plausible because increased cell-to-cell variability in specific phenotypes (in our case, cell length and division time) also increases the number of cells with a sub-optimal phenotype. In an unchanging or optimal environment, such maladapted cells decrease the mean trait value, leading to reduced population growth rate (Fig. 8). Indeed, in fluctuating or stressful environments, both predictions (Zhuravel et al. 2010) and prior observations (Levy et al. 2012) suggest that cell to cell variability can be beneficial, and hence more likely to persist. Thus, it is possible that under fluctuating environments, cell-to-cell variation leads to increased variance across population fates. Further work – including modelling efforts – may help to distinguish between these possibilities, and resolve conditions under which single-cell (between-individual) heterogeneity should also increase between-population variance.

Our results also indicate that the underlying cause of mistranslation may play a large role in determining the impact of mistranslation-induced variability. The effect of mistranslation varied depending on the mechanism used to increase mistranslation, as well as across different population growth parameters (growth rate, yield and lag time). These inconsistencies could stem from multiple factors: i) Each environmental manipulation (canavanine, norleucine, streptomycin) affects different sets of proteins. For example, canavanine and norleucine should respectively alter proteins rich in arginine and methionine; while streptomycin has a global effect, impacting all protein production. Note that in addition to increasing variability in population parameters, streptomycin treatment also shows a trend towards increasing variability in all population parameters measured (Fig. 6f). ii) Secondary impacts unrelated to mistranslation may complicate the relationship between the degree of mistranslation and phenotypic variability. For example, norleucine also inhibits DNA methylation and methionine biosynthesis (Bogosian et al. 1989). To generate hyper-accurate ribosomes, we used a mutation that impacts population growth rate minimally (Samhita et al. 2020), and the WT single cell data show tighter distributions for both division time and cell length as compared with their parent strains (Fig. 2b–c, Table S1). However, lag time across population replicates was more variable than in the parent strain both in the WT and Mutant (Fig. 3e). Although we do not have a good explanation for this, the combination of two mutations in the Mutant background could lead to other epistatic effects. Also, hyper-accurate ribosomes are expected to drop off more often during translation (Karimi and Ehrenberg 1994), potentially increasing variability in the number of actively dividing cells at any given time, and altering lag time. Given these complex relationships, it is perhaps not surprising that we do not see a consistent trend linking various causes of mistranslation with phenotypic variability. Thus, our work identifies the mechanistic basis of mistranslation as an important factor to understand the phenotypic effects of mistranslation.

Our work addresses gaps in prior work, broadening our understanding of the potential role of mistranslation in evolutionary dynamics. Previous studies found that mistranslation is correlated with variability in phenotype (Bacher et al. 2007; Bezerra et al. 2013) but could not establish a causal link. Others identified specific mechanisms that increased stress resistance under global mistranslation mediated by altered proteomes (Fan et al. 2015), but did not establish whether this relied on a general increase in phenotypic variability. In our work, it is reasonable to assume that a “statistical proteome” is generated in each mutant cell as a result of mistranslation (Winther and Gerdes 2011; Samhita et al. 2013). Depleting initiator tRNA content (as in our mutant) can alter the cellular proteome in various ways: via non AUG initiation (Winther and Gerdes 2011; Samhita et al. 2013), ribosome alterations (Shetty and Varshney 2016) or by simply lowering translation rates (Samhita et al. 2014), potentially leading to instantaneous changes in global transcript as well as protein levels. We show that completely different mechanisms of mistranslation (non-AUG initiation and decreased translation; replacing an amino acid with a non-native analogue; increasing decoding errors) all converge on the pattern of increased variability in growth and survival. Thus, the potentially varied and specific mechanisms linking various forms of mistranslation to phenotypic variation ultimately achieve the same end point under stress: an increased probability of population survival. In addition, compared to prior work, our results are somewhat more applicable to *E. coli* function and evolution in natural ecosystems. For instance, the ecological relevance of genetic manipulations used in prior work (such as mutations in the ribosomal protein S4 (Fan et al. 2015; Bratulic et al. 2017)) is unclear. In contrast, translation – and specifically translation initiation – is reduced in response to several environmental stresses (Nagase et al. 1988; Winther and Gerdes 2011; Watanabe et al. 2013), creating a cellular environment analogous to that of our mistranslating mutant with reduced initiator tRNA. Second, we used two stressful conditions – starvation and high temperature – that lead to increased cell death in *E.coli* and are thought to be encountered by *E.coli* in its natural habitat (reviewed in Koch 1971; van Elsas et al. 2011). Finally, we measured the impact of mistranslation on cell growth and survival, which are key parameters governing microbial ecological and evolutionary dynamics and have significant repercussions for genome structure and evolution (Roller et al. 2016). Thus, we speculate that cells could modulate mistranslation levels as a generalized, global response to multiple stressful conditions that they encounter in nature.

In closing, we note that our results lead to a number of interesting open questions. For instance, while we observe that mistranslation reliably generates variability, we do not know if the same variants are re-generated across generations; precisely which variants survive under each stressful condition; and whether this is predictable across environments. More work is also needed to clarify whether advantageous variants pave the way for the phenotype to be fixed by mutation, as suggested previously (Cowen and Lindquist 2005; Whitehead et al. 2008). Finally, while our experiments inform about the short-term impact of altering mistranslation, the longer-term impacts of mistranslation need to be investigated further. For instance, recent work showed that a mistranslating strain fixes distinct sets of beneficial mutations during laboratory adaptation to antibiotic stress (Bratulic et al. 2017). It would be exciting if these results could be generalized across various stresses and forms of mistranslation. Here, we have built a foundation to address these questions by demonstrating that mistranslation can influence short-term population trajectories and set the stage for longer-term evolutionary consequences.

## MATERIALS AND METHODS

### Bacterial strains

To manipulate mistranslation levels in wild type (WT) KL16 *E. coli* cells (Low 1968), we used two genetically altered derivatives of the WT. As our focal ‘mistranslating’ strain, we used the KLΔ*ZWV* strain (henceforth ‘mutant’), which lacks three of the four initiator tRNA genes encoded by *E. coli* (Samhita et al. 2013). Initiator tRNA acts only at the first step of protein synthesis and has no substitute (Gualerzi and Pon 2015). Since the mutant carries ∼25% of the WT initiator tRNA complement, it has a lower rate of protein synthesis, a ∼20% slower growth rate than the WT (Samhita et al. 2020), and mistranslates through non-AUG initiation (Samhita et al. 2013). In contrast, to reduce mistranslation rates in the WT, we introduced a mutation (K42R) in the protein S12 that increases translation accuracy by reducing the frequency of decoding errors (Chumpolkulwong et al. 2004). We transferred this mutation into KL16 and KLΔ*ZWV* from the parent strain SS3242 obtained from CGSC, Yale university, via P1 transduction, generating strains KL(HA) and KLΔ*ZWV*(HA), referred to as WT(HA) and Mutant(HA) in the text. The mutation led to an ∼10-fold increase in translation accuracy both in the WT and in the Mutant (Samhita et al. 2020). For single-cell variability measurements, we used WT and mutant strains carrying a genomically encoded, constitutively expressed GFPmut2 allele tagged with a kanamycin resistance marker inserted between the genes *aidB* and *yjfN*, and expressed from a P5 promoter (gifted by Prof Bianca Sclavi, ENS, Paris).

### Growth conditions and media

When generating strains or to simulate control (normal growth) conditions, we grew bacterial cultures in Luria Bertani medium (LB) or on LB-agar plates containing 1.8% (w/v) agar (Difco), incubated at 37°C. In some experiments, we also altered growth conditions to elevate mistranslation levels and/or impose stress, as follows. To increase mistranslation levels, we added (1) canavanine sulphate at concentrations ranging from to 0.375 to 3 mg/mL as specified in each experiment (canavanine is an analogue of arginine that induces mistranslation) (Fan et al. 2015), (2) norleucine (an analog of leucine which substitutes for the amino acid methionine in proteins (Karkhanis et al. 2007)) at concentrations ranging from 0.28 to 2.25 μg/mL and (3) streptomycin sulphate at concentrations ranging from 0.625 to 5 μg/mL (streptomycin leads to errors in ribosomal decoding (Carter et al. 2000)). To impose stress, we subjected single cells to starvation by supplying only saline instead of a growth medium (i.e. no nutrients), or by stopping the flow of growth media, allowing entry into the stationary phase of growth. At the population level, we let cultures grow till late stationary phase when nutrients are depleted, which imposes stress on the cells (Koch 1971). Finally, we cultured cells in LB at high temperature (42°C), imposing stress that reduces survival (van Elsas et al. 2011). To check the impact of a transient increase in mistranslation on survival at 42°C, we exposed replicate cultures to different concentrations of canavanine, norleucine or streptomycin for ∼2 h until they reached OD_600_ ∼0.6. We then pelleted and re-suspended cells and cultured them for 2 h at 42°C before carrying out dilution plating to assess survival. Each treatment had triplicates before and after exposure to 42°C, i.e. a total of 6 cultures. Given three concentrations and three mistranslating agents, this led to 18*3= 54 agar plates, which were done in staggered sets of 18 each. Handling more than 18 at a time led to significant time variation in the steps of cell pelleting and exposure to 42°C, adding to the variability that we were attempting to capture. As a result, at any given time, we were limited to three treatments for a given mistranslating agent. In addition, we could not compare across multiple blocks of the same experiment because of the high across-experiment variability in growth and colony numbers.

### Single cell microfluidics measurements

To measure variability in growth characteristics at the single cell level, we used a microfluidic device seeded with GFP-labelled *E. coli* strains (described above). The ‘mother machine’ microfluidic device was fabricated as previously described (Wang et al. 2010). Saturated overnight cultures of WT, Mutant, WT(HA) and Mutant(HA) were sub-cultured to 1% by volume into LB and incubated at 37 C for 3 h. To concentrate cells ∼20 fold, we centrifuged the cultures (5 min, 3000g) and re-suspended cells in 200 μL LB. We injected the cell suspension into the microfluidic device using a syringe, and allowed the cells to diffuse into the growth channels (∼ 2 h; see schematic in Fig. 4a). Then, the device was placed in a temperature-controlled stage-top incubator (OKOlab), which in turn was placed on an inverted microscope (Olympus IX81). To allow cell growth in the device, we pumped LB from a 15 mL centrifuge tube (Greiner) held at constant temperature in a dry block heater (IKA). We allowed cells to grow in the device for 2.5 h at a media flow rate between 600 – 800 μL/h, and then began imaging cells to measure growth characteristics. When necessary, stationary phase growth was simulated by stopping media inflow after 190 min of image acquisition, and then allowing cells ∼3 h (∼6 doublings) to enter stationary phase before readings were taken. We used a coolLED lamp (excitation: 490 nm) for fluorophore excitation, and captured bright field and fluorescence images at intervals of 2 min for 10 h at 40X magnification using an EMCCD camera (Photometrics Prime). We imaged ∼160 channels at constant flow and temperature and carried out preliminary image editing using ImageJ and used a custom MATLAB code (MathWorks) to extract information on cell length and division time. We cut individual channels in ImageJ and used a custom MATLAB code (MathWorks) to binarize the images and assign an identity to each cell. Based on changes in fluorescence intensity in the cell body, the code identified a cell division. We measured cell length at every frame to calculate cell length at birth and division, and the corresponding time for each event. When counting live and dead cells under stress, an instantaneous loss of fluorescence signal was used as an indicator of cell death.

### Measuring population growth and yield

To measure variability in growth across populations, we used 40 independent colonies of each *E. coli* strain as biological replicates. We inoculated colonies in LB and allowed them to grow overnight at 37°C with shaking at 200 rpm for 16 hours. We then added 5 μL of the overnight culture into 495 μL of the relevant growth medium in 48 well microplates (Corning-Costar), and incubated them in a shaking tower (Liconic) at 37°C. We measured optical density (OD) of each well at 600 nm using an automated growth measurement system (F100-Pro, which includes a microplate reader from Tecan, Austria), every 30 or 40 minutes for 12 to 18 hours. The automated system allowed us to simultaneously measure growth rates in up to 10 microplates. We estimated maximum growth rate using the Curve Fitter software (Delaney et al, 2013) and maximum OD value (OD_max_, as a proxy for growth yield) by averaging the five highest OD values.

### Measuring cell survival

We measured cell survival in liquid culture by plating serial dilutions of the culture on agar medium and counting colonies (which represent viable cells from the original culture). Briefly, we set up 20 replicate cultures of a strain (each inoculated from an individual colony) and allowed cultures to grow to saturation overnight in rich media (LB) or under stress, as required. We then sub-cultured cells 1% by volume and set up the experiment. At appropriate time intervals, we serially diluted the culture until we obtained sufficiently dilute cultures such that the final dilution plated on LB agar would give rise to ∼ 20 to 200 distinct colonies (which can be counted reliably). We used 55 μL of this focal culture, diluted it in 495 μL of normal saline (to generate a 1:10 dilution), and vortexed the mix thoroughly. We continued diluting serially until we obtained sufficiently dilute cultures such that 100 μLμL of the culture plated on LB agar would give rise to ∼ 20 to 200 distinct colonies (which can be counted reliably). We incubated plates for 24 hours, counted colonies and multiplied by the appropriate dilution factor to determine viable counts in the original culture.

### Measuring competitive fitness

To test the relative fitness of WT and mutant strains under competition, we first had to establish a way to distinguish the two strains. To do so, we generated KLΔ*lacZ* (WT lacking the *lacZ* gene). This strain forms white colonies on MacConkey’s agar, while the other strains under study – Mutant, WT(HA) and Mutant(HA) – form pink colonies because they carry an intact *lacZ* gene. We grew the two strains being competed to saturation (overnight) and then mixed them (1% each by volume) into 5 mL of growth medium. We then subjected the mix to periodic dilution plating onto MacConkey’s agar, and determined the relative numbers of each strain over time. We confirmed that the *lacZ* deletion is selectively neutral, by competing it against the unmarked parent WT strain (Fig. S1).

### Quantifying variability

As described above, we collected data on the size, length, and division time of single cells; and parameters such as growth rate, growth yield (OD_max_) and lag time for replicate populations of each strain or experimental treatment. In most cases, the data were not distributed normally (Shapiro Wilke test for normality). Therefore, we could not use standard quantifications of variability such as the variance or coefficient of variation (CV=standard deviation/mean). To assess differences in variability across groups, we employed a non-parametric statistical test-Fligner-Killeen test-that is robust to departures from normality (Fligner and Killeen 1976; Conover et al. 1981). This test ranks all data around the median value, measures the values of the residuals for each data point, and calculates the test statistic by ranking the residual values. All comparisons are tabulated in Table S1. As appropriate, we visualized variability using the frequency distribution of each measured variable, or the inter quartile range of the data.

## ACKNOWLEDGEMENTS

We thank Vidyanand Nanjundiah and members of the Agashe lab for discussion and critical comments on the manuscript, and Debshankar Banerjee for the MATLAB code used for analysing the single cell data. We acknowledge funding and support from the Wellcome Trust/DBT India Alliance (grant IA/E/14/1/501771 to LS and grant IA/I/17/1/503091 to DA); the Indian Council for Medical Research (3/1/3/JRF-2015 to PR); the Simons Foundation (ST and GS); the Max Planck Society through a Max Planck Partner Group (ST); the Council for Scientific and Industrial research (CSIR research grant 37(1629)/14/EMR-II to DA); and the National Centre for Biological Sciences (NCBS-TIFR).

## AUTHOR CONTRIBUTIONS

LS and DA conceived the project; LS, DA and ST designed experiments; LS, PR and GS conducted experiments; LS, GS and DA analysed data; LS and DA wrote the manuscript with input from all authors.

