## Supplementary material for "THE IMPACT OF MISTRANSLATION ON PHENOTYPIC VARIABILITY AND FITNESS"

| Figure details | Comparison | Test | Median values | Statistic | P value |
|---|---|---|---|---|---|
| **Fig.2 (b-e)** | | | | | |
| Time to division (minutes), 37°C | WT vs Mutant | Mann-Whitney | 36 vs 62 | U=45832 | P<0.0001 |
| | WT vs WT(HA) | Mann-Whitney | 36 vs 38 | U=129812 | P=0.5667 |
| | Mutant vs Mutant (HA) | Mann-Whitney | 62 vs 56 | U=109351 | P<0.0001 |
| | WT vs $WT_{can}$ | Mann-Whitney | 36 vs 80 | U=20542 | P<0.0001 |
| | WT vs $WT_{nor}$ | Mann-Whitney | 36 vs 48 | U=79978 | P<0.0001 |
| | WT vs $WT_{strp}$ | Mann-Whitney | 36 vs 44 | U=103802 | P<0.0001 |
| Length at birth (micron), 37°C | WT vs Mutant | Mann-Whitney | 2.3 vs 2.7 | U=584346 | P<0.0001 |
| | WT vs WT(HA) | Mann-Whitney | 2.4 vs 2.2 | U=708782 | P<0.001 |
| | Mutant vs Mutant (HA) | Mann-Whitney | 2.7 vs 2.9 | U=664722 | P<0.0001 |
| | WT vs $WT_{can}$ | Mann-Whitney | 2.36 vs 2.25 | U=749113 | P<0.0001 |
| | WT vs $WT_{nor}$ | Mann-Whitney | 2.36 vs 2.7 | U=497861 | P<0.0001 |
| | WT vs $WT_{strp}$ | Mann-Whitney | 2.36 vs 2.34 | U=939394 | P=0.36 |
| **Variability comparisons:** | | | **Difference in variability (Yes/No)** | | |
| Time to division (minutes), 37°C | WT vs Mutant | Fligner | Yes | $X^2$=81.8 | P=2.2x$10^{-16}$ |
| | WT vs WT(HA) | Fligner | Yes | $X^2$=5.3 | P=0.02 |
| | Mutant vs Mutant (HA) | Fligner | No | $X^2$=1.4 | P=0.2 |
| | WT vs $WT_{can}$ | Fligner | Yes | $X^2$=87.6 | P=2.2x$10^{-16}$ |
| | WT vs $WT_{nor}$ | Fligner | Yes | $X^2$=66.9 | P=1.2x$10^{-13}$ |
| | WT vs $WT_{strp}$ | Fligner | Yes | $X^2$=32.9 | P=1.18x$10^{-8}$ |
| Length at birth (micron), 37°C | WT vs Mutant | Fligner | Yes | $X^2$=91.4 | P=2.2x$10^{-16}$ |
| | WT vs WT(HA) | Fligner | Yes | $X^2$=40.4 | P=5.75x$10^{-8}$ |
| | Mutant vs Mutant (HA) | Fligner | Yes, Mutant(HA)>Mutant | $X^2$=50.5 | P=1.17x$10^{-12}$ |
| | WT vs $WT_{can}$ | Fligner | Yes, $WT_{can}$>WT | $X^2$=49.0 | P=2.5x$10^{-12}$ |
| | WT vs $WT_{nor}$ | Fligner | Yes | $X^2$=58.5 | P=2.03x$10^{-14}$ |
| | WT vs $WT_{strp}$ | Fligner | No | $X^2$=1.5 | P=0.2 |
| **Fig. 3** | | | | | |
| **Variability comparisons:** | | | **Difference in variability (Yes/No)** | | |
| Population growth rates | Mutant vs WT | Fligner | No | $X^2$=0.0009 | P=0.9 |
| | WT(HA) vs WT | Fligner | No | $X^2$=1.3 | P=0.2 |
| | Mutant(HA) vs Mutant | Fligner | Yes | $X^2$=5.8 | P=0.02 |
| | WT(can) vs WT | Fligner | No | $X^2$=1.7 | P=0.2 |
| | WT(nor) vs WT | Fligner | No | $X^2$=1.2 | P=0.26 |
| | WT(Strp) vs WT | Fligner | Yes | $X^2$=10.6 | P=0.001 |
| Population growth yields | Mutant vs WT | Fligner | No | $X^2$= 3 | P=0.08 |
| | WT(HA) vs WT | Fligner | No | $X^2$=1.1 | P=0.28 |
| | Mutant(HA) vs Mutant | Fligner | No | $X^2$=2.5 | P=0.1 |
| | WT(can) vs WT | Fligner | No | $X^2$=0.8 | P=0.3 |
| | WT(nor) vs WT | Fligner | Yes | $X^2$=8.1 | P=0.004 |
| | WT(Strp) vs WT | Fligner | Yes | $X^2$=31.3 | P=2.18x$10^{-8}$ |

| | | | | | |
|---|---|---|---|---|---|
| Population lag times | Mutant vs WT | Fligner | No | $X^2$= 3 | P=0.08 |
| | WT(HA) vs WT | Fligner | Yes | $X^2$=5.3 | P=0.02 |
| | Mutant(HA) vs Mutant | Fligner | Yes | $X^2$=37.4 | P=9.24x$10^{-10}$ |
| | WT(can) vs WT | Fligner | No | $X^2$=3.5 | P=0.06 |
| | WT(nor) vs WT | Fligner | No | $X^2$=0.4 | P=0.5 |
| | WT(Strp) vs WT | Fligner | Yes | $X^2$=13.7 | P=0.0002 |
| **Fig. S4** | | | | | |
| Population growth rate median comparisons | Mutant vs WT | Mann-Whitney U | 0.9 vs 1.15 | U= 4 | P=0.08 |
| | WT(HA) vs WT | Mann-Whitney U | 0.79 vs 1.15 | U=0 | P<0.0001 |
| | Mutant(HA) vs Mutant | Mann-Whitney U | 0.96 vs 0.9 | U=586 | P=0.008 |
| | WT(can) vs WT | Mann-Whitney U | 0.93 vs 1.15 | U=0 | P<0.001 |
| | WT(nor) vs WT | Mann-Whitney U | 1.17 vs 1.15 | U=729 | P=0.04 |
| | WT(Strp) vs WT | Mann-Whitney U | 1.1 vs 1.15 | U=504 | P=0.0006 |
| Population growth yield median comparisons | Mutant vs WT | Mann-Whitney U | 0.8 vs 0.76 | U=286 | P<0.0001 |
| | WT(HA) vs WT | Mann-Whitney U | 0.73 vs 0.8 | U=159 | P<0.0001 |
| | Mutant(HA) vs Mutant | Mann-Whitney U | 0.88 vs 0.73 | U=0 | P<0.0001 |
| | WT(can) vs WT | Mann-Whitney U | 0.86 vs 0.8 | U=253 | P<0.001 |
| | WT(nor) vs WT | Mann-Whitney U | 0.93 vs 0.8 | U=0 | P=0.04 |
| | WT(Strp) vs WT | Mann-Whitney U | 0.54 vs 0.8 | U=6 | P<0.0001 |
| Population lag time median comparisons | Mutant vs WT | Mann-Whitney U | 1.26 vs 1 | U=78 | P<0.0001 |
| | WT(HA) vs WT | Mann-Whitney U | 0.48 vs 1 | U=14 | P<0.0001 |
| | Mutant(HA) vs Mutant | Mann-Whitney U | 0.98 vs 1.26 | U=497 | P=0.002 |
| | WT(can) vs WT | Mann-Whitney U | 1.6 vs 1 | U=0 | P<0.0001 |
| | WT(nor) vs WT | Mann-Whitney U | 0.68 vs 1 | U=0 | P<0.0001 |
| | WT(Strp) vs WT | Mann-Whitney U | 1.4 vs 1 | U=128 | P<0.0001 |
| **Fig.4a** | | | | | |
| Time to division (minutes), 42°C | WT vs Mutant | Mann-Whitney | 28 vs 38 | U=123307 | P<0.0001 |
| | Mutant vs Mutant(HA) | Mann-Whitney | 38 vs 40 | U=149832 | P=0.0031 |
| | WT vs WT(HA) | Mann-Whitney | 28 vs 34 | U=118457 | P<0.0001 |
| **Fig. 7a** | | | | | |
| | WT (can -3) vs WT | Mann-Whitney U | 48.5 vs 11 | U=0 | P<0.0001 |
| | WT (nor-2.25) vs WT | Mann-Whitney U | 32 vs 11 | U=0 | P<0.001 |
| | WT (strp-5) vs WT | Mann-Whitney U | 39 vs 11 | U=0 | P<0.0001 |
| **Fig. 7b** | | | **Means** | | |
| | WT (can-3) vs WT | Paired t test | 13.5 vs 1.7 | t=0.96 | P=0.4 |
| | WT (nor 2.25) vs WT | Paired t test | 1.25 vs 1.7 | t=0.75 | P=0.5 |
| | WT (strp 5) vs WT | Paired t test | 30 vs 1.7 | t=2.9 | P=0.1 |
