## Supplementary figures for "THE IMPACT OF MISTRANSLATION ON PHENOTYPIC VARIABILITY AND FITNESS"

Figure S1

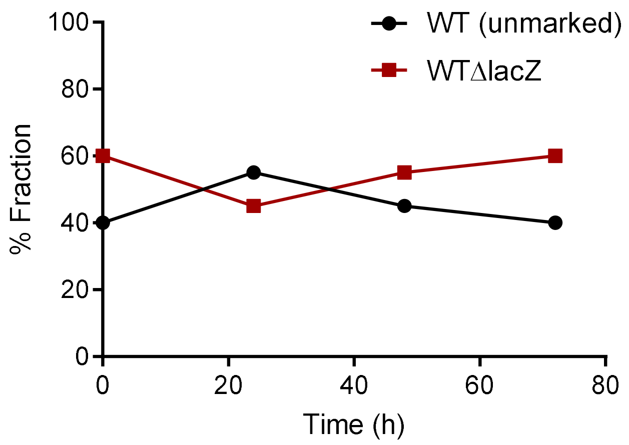


**Figure S1. *lacZ* knock out does not influence the outcome of pair-wise competition experiments:** Log phase cultures ($OD_{600}$~0.6) of WT and WTΔ*lacZ* were subjected to pair-wise competition experiments in LB at 37°C followed by plating on MacConkey's agar. Percentage fraction of each strain is plotted against time here.

Figure S2

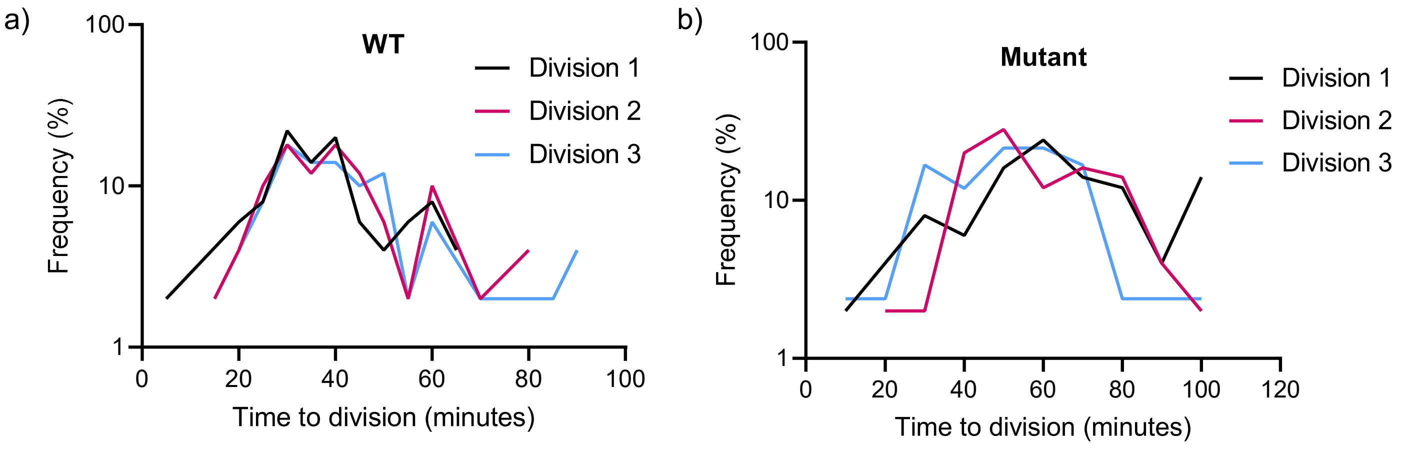


**Figure S2. Distributions of time to division for individual mother cell divisions:** Individual mother cells (n=3) from WT and Mutant were monitored in the microfluidics device for ~60 cell divisions each, at 37$^{o}$C. Frequency distributions arising from each such lineage of divisions is plotted here against time to division. For WT, comparison of means across the three distributions, Welch's ANOVA test, W=1.9, P=0.15. For Mutant, comparison of means across the three distributions, Welch's ANOVA test, W=4.3, P=0.02

Figure S3

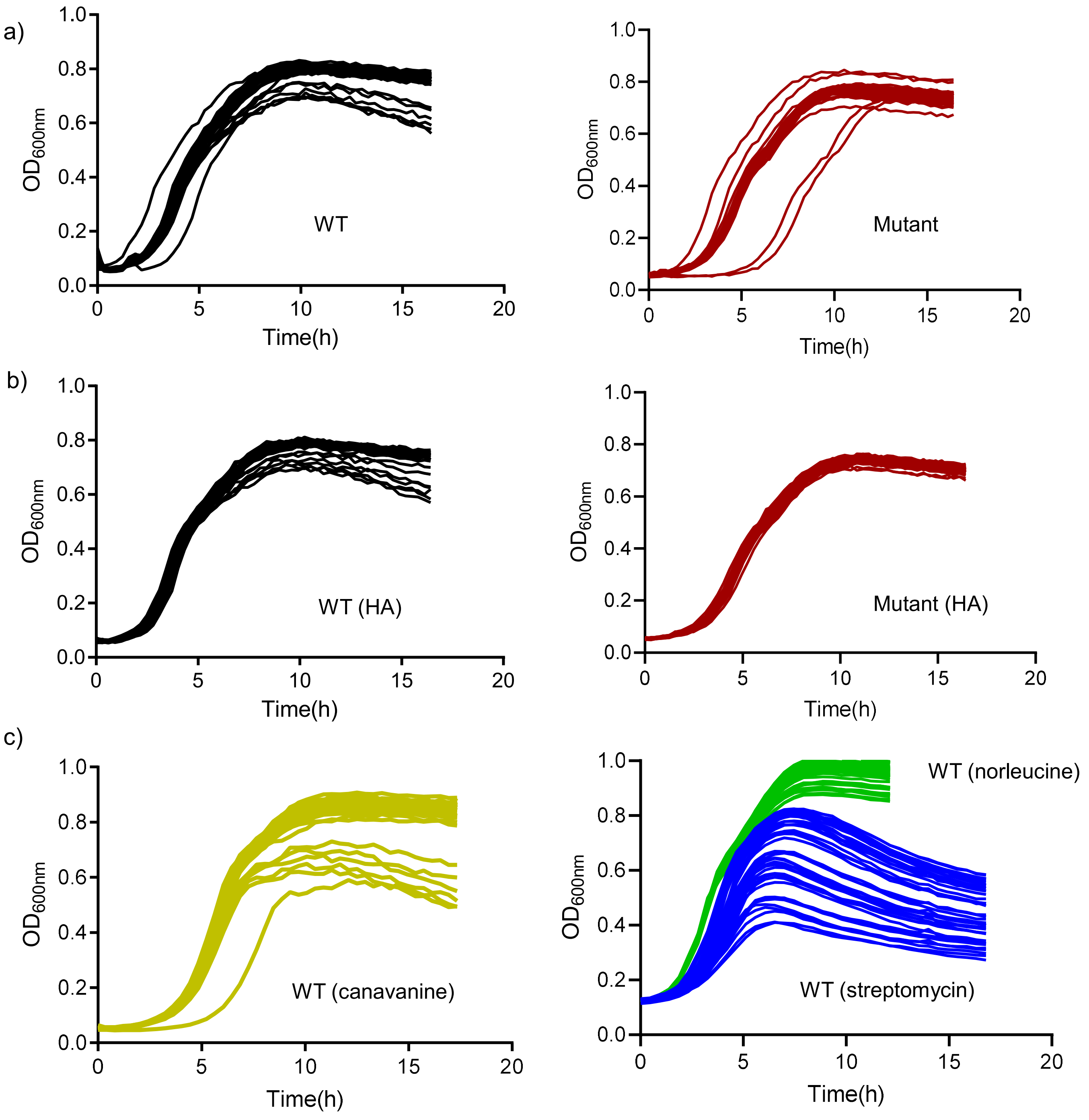


**Figure S3. Raw growth curves showing independent biological replicates:** Raw growth curves for the strains indicated (n=40) showing $OD_{600}$ over time plots for each biological replicate (corresponding to a single colony) as obtained by a Tecan growth reader recording $OD_{600}$ every 30 minutes. Canavanine (3 mg/mL), Norleucine (2.25 μg/mL) and Streptomycin (5 μg/mL) were added to the growth medium of LB where indicated.

Figure S4

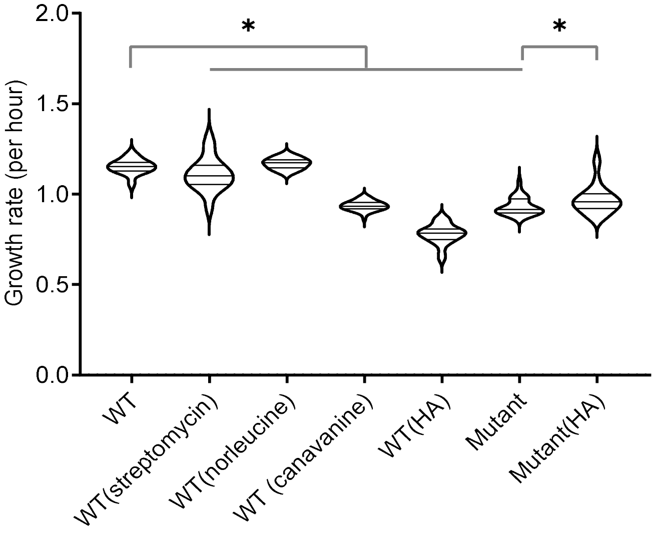


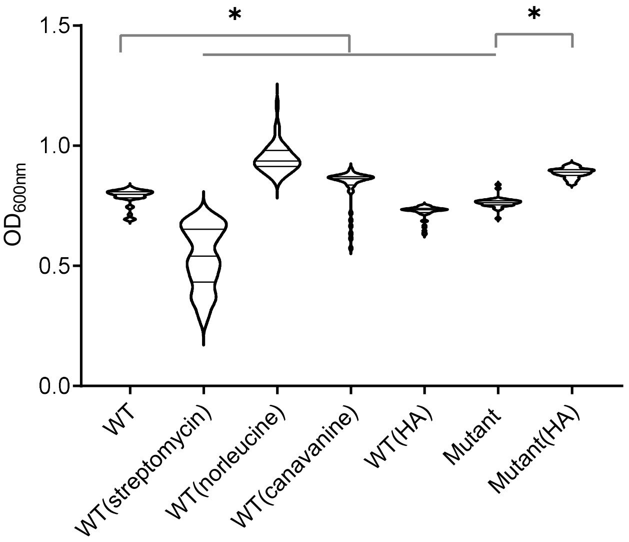


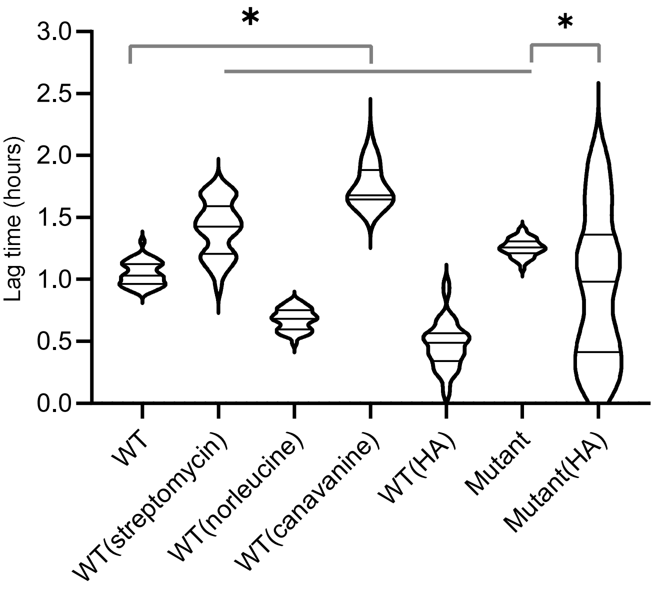


**Figure S4. Mistranslation impacts mean fitness across replicate populations:** Violin plots showing distributions of three population growth parameters, estimated using ~40 (37 to 44) biological replicates (populations) for each strain or growth condition. (a–b) Growth rate (c–d) growth yield and (e–f) lag time (time until culture reaches $OD_{600}$ ~0.02). Median, $25^{th}$ and $75^{th}$ quartiles are indicated by solid lines within each violin. The length of each violin corresponds to the range of the distribution. Asterisks indicate significant differences in median values. WT=wild type; HA=hyper-accurate.

Figure S5

| | Set 1 | | Set 2 | | Set 3 | |
|---|---|---|---|---|---|---|
| | Live | Dead | Live | Dead | Live | Dead |
| WT | 160 | 37 | 286 | 73 | 329 | 285 |
| Mutant | 197 | 10 | 409 | 19 | 69 | 9 |

Figure S6

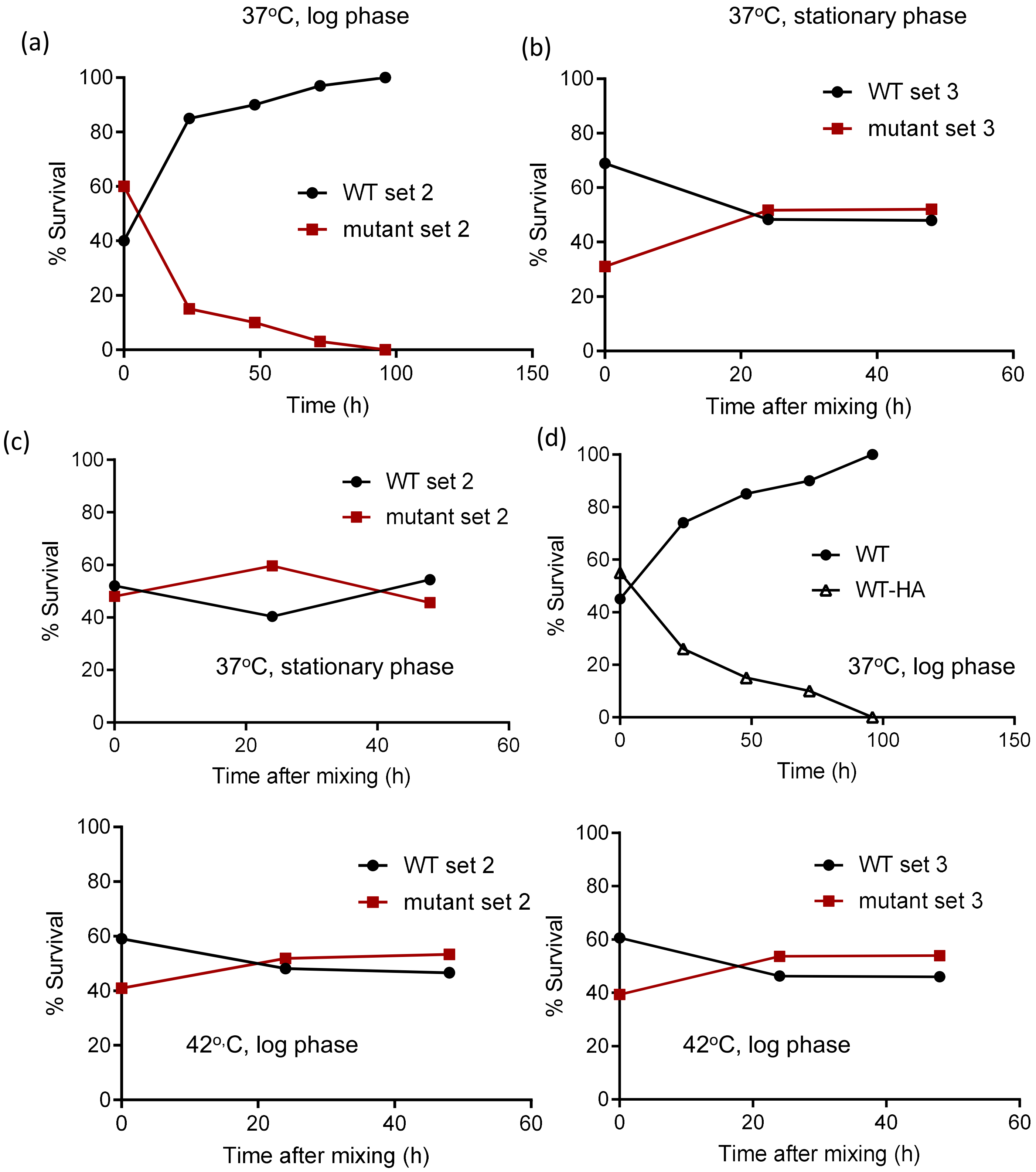


**Figure S6. Replicate blocks for pair-wise competition experiments:** Pair-wise growth competition experiments, strains and conditions as indicated, see Fig.4b-e.
